# Modeling herpes simplex virus type 2 and Zika virus replication in vaginal organoids and spheroids

**DOI:** 10.64898/2026.03.02.709097

**Authors:** Ananya Saraph, Imogen Porter, Grace Bodykevich, Ingrid Barta, May Dang-Lawson, Maria Tokuyama

## Abstract

Sexually transmitted viral infections disproportionately affect women and have adverse outcomes for maternal and fetal health. Since they are acquired through the vaginal tract, the vaginal epithelium is an important site to study host-viral interactions. An *in vitro* model of mouse vaginal epithelial organoids that can be infected by herpes simplex virus was recently established. Here, we used multiple vaginal organoid systems and studied their response to infection by herpes simplex virus type 2 (HSV-2) and Zika virus (ZIKV), both of which can be sexually transmitted and result in adverse effects on maternal and fetal health. We showed that mouse vaginal organoids support the replication of HSV-2 and ZIKV and respond to interferon treatment. We also established human vaginal spheroids from VK2/E6E7 cells that contain distinct apical and basal cell layers and are also susceptible to HSV-2 and ZIKV infection. Acyclovir treatment reduced HSV-2 replication in both mouse organoids and VK2 spheroids. However, ZIKV was restricted by remdesivir in VK2 spheroids, but not in mouse organoids, indicating differences in antiviral activity depending on the organoid system. Finally, we established apical-out mouse vaginal organoids that were also susceptible to HSV-2 and ZIKV infection. Altogether, our results show that vaginal organoids are useful *in vitro* models to study vaginal viral infections and effects of antiviral drugs.

## Introduction

Sexually transmitted viral infections, including human papillomavirus (HPV), human immunodeficiency virus (HIV), herpes simplex virus (HSV), and Zika virus (ZIKV) are a significant global health concern due to their high morbidity and adverse impacts on reproductive health (1). These sexually transmitted viruses (STVs) disproportionately affect women, often resulting in adverse outcomes for maternal, fetal, and neonatal health (2–7). With the exception of HPV, no vaccines are available to prevent these infections (8, 9) and antivirals are limited (10–12). Therefore, better understanding of the virus-host interaction for STVs is needed.

The vaginal tract is the main route of infection by STVs in women. It is part of the lower female reproductive tract (FRT) and has a type II mucosal surface lined with a stratified squamous epithelium, in contrast to the simple columnar epithelium of the upper FRT (13). Many *in vivo* and *in vitro* models are used to study the development and pathologies of the FRT, including *in vitro* models of the FRT in the form of three-dimensional (3D) organoids (14). For example, organoids of the fallopian tubes, ovaries, cervix, placenta and endometrium are used to study conditions such as cervical carcinogenesis, HPV infection, endometriosis, and placental formation (15–20). Recent studies showed successful establishment of vaginal organoids from mouse tissues and their susceptibility to HSV infection (21, 22). However, the use of vaginal organoids is still limited, and there are no organoid models of the human vaginal tract.

The vaginal organoids derived from stem cells in the mouse vaginal tract are self-renewing and contain distinct apical and basal surfaces as observed in the vaginal epithelium (21). These organoids display a basal-out orientation such that the basal cells are exposed to the external environment, in contrast to the typical epithelial structure wherein apical cells are external facing. Despite the flipped orientation, these organoids support tissue-resident memory CD8^+^ T cells and are susceptible to HSV-1 and HSV-2 infection (22), suggesting that vaginal organoids can be used to study vaginal infections and inform novel host-pathogen interactions. However, many aspects of vaginal organoids remain undetermined, including susceptibility to other viruses, immune responses to infection, and effects of antiviral drugs. The implications of the basal-out orientation on viral pathogenesis and host response are also not clear.

In order to investigate these characteristics, we generated both basal-out and apical-out vaginal organoids from mouse tissues, as well as vaginal spheroids from the transformed human vaginal epithelial cell line, VK2/E6E7. All three systems supported HSV-2 and ZIKV replication. Mouse vaginal organoids were able to mount an interferon (IFN) response to HSV-2 infection. HSV-2 and ZIKV replication were restricted upon treatment with acyclovir and remdesivir, respectively. Together, our work revealed features of the vaginal organoids that can help inform future studies on host-pathogen interactions using these models.

## Results

### Characterization of primary mouse vaginal organoids and human VK2 spheroids

To study vaginal viral infections in primary mouse vaginal organoids, we obtained whole vaginal tissues from mice treated with depot medroxyprogesterone acetate (DMPA), which synchronizes mice in a diestrus-like state and increases susceptibility to vaginal infections including HSV-2, HIV, and ZIKV (23–25) (Fig. 1A). We digested these tissues to release vaginal stem cells that are capable of forming self-renewing organoids that mimic the stratified structure of the squamous epithelium (21). We then cultured total cells in the presence of Rho-associated coiled-kinase (ROCK) inhibitor to reduce apoptosis, transforming growth factor beta (TGFβ) receptor kinase inhibitor to stimulate cellular differentiation, and epidermal growth factor (EGF) to promote growth and maintain the stem cell-like nature of vaginal cells. These cells developed into organoids that progressively grew in size and formed distinct basal and suprabasal layers by day 7 post-culture (Fig. 1B).

**Fig 1.**
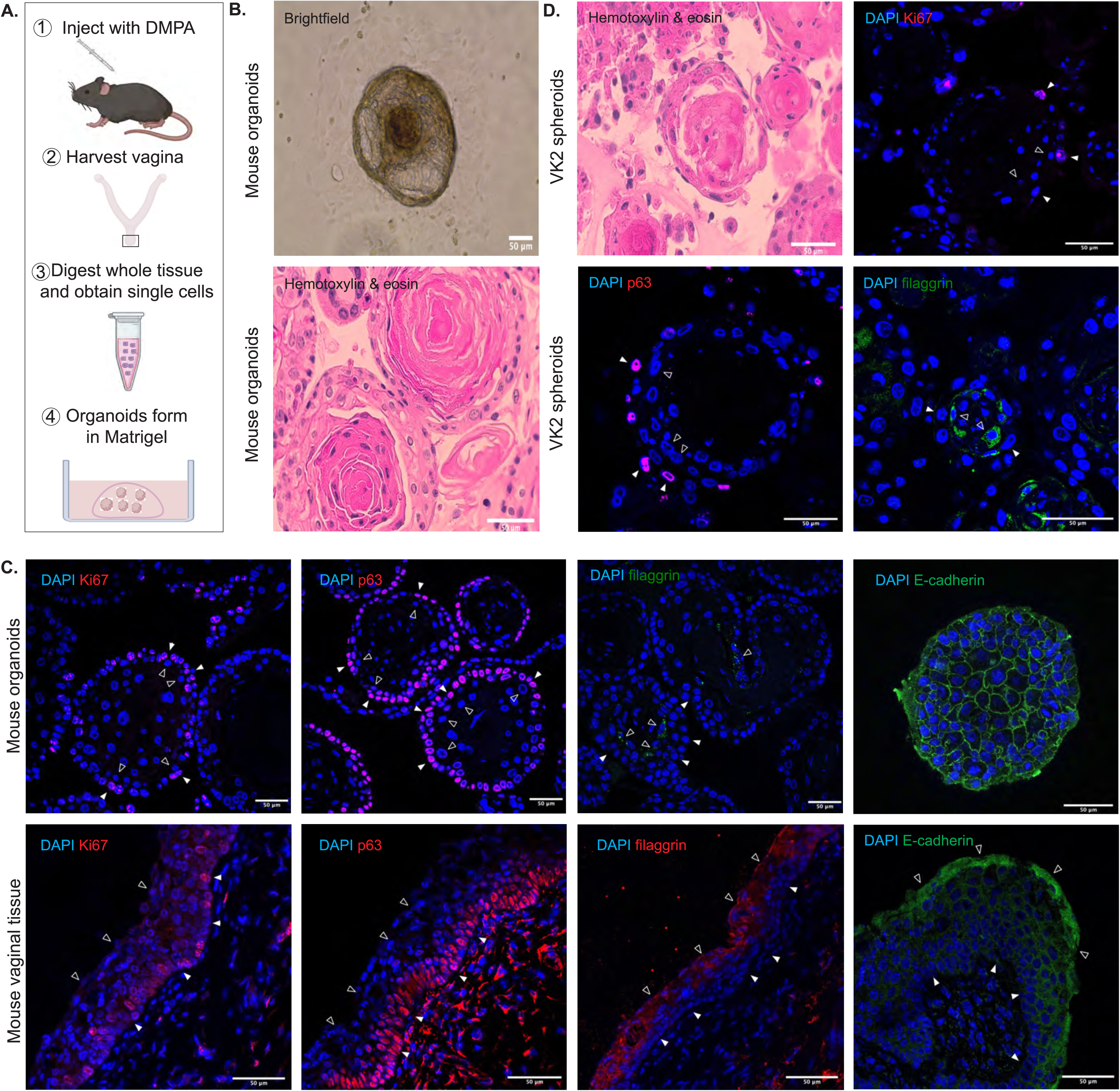
Establishment and characterization of mouse vaginal organoids and VK2 spheroids. (A) A schematic illustration of the establishment of mouse vaginal organoids. (B) Brightfield image of 7-day old mouse vaginal organoids (top) and H&E staining of 10-day old mouse vaginal organoid sections (bottom). (C) Sections of mouse vaginal organoids (top) and mouse vaginal tissue (bottom) stained for Ki67, p63, Filaggrin, and E-cadherin. (D) Sections of VK2 spheroids stained for H&E, Ki67, p63, and filaggrin. Magnification 20X (brightfield) and 63X; scale bars represent 50 µm. Solid arrows indicate basal cells, and open arrows indicate apical cells.

Hematoxylin and Eosin (H&E) staining of vaginal organoid sections revealed distinct basal and apical layers (Fig. 1B). Distinct epithelial layers were detected between days 10 and 14, as indicated by markers of cell proliferation (Ki67), basal cells (p63), apical cells (filaggrin), and cell to cell adhesion (E-cadherin) (Fig. 1C, top). The outer layer consisted of proliferating Ki67+ cells that expressed p63, while cells in the inner layer expressed filaggrin, consistent with the basal-out orientation. E-cadherin was expressed at cell-to-cell junctions throughout the organoids. These expression patterns were comparable to that of vaginal tissues (Fig. 1C, bottom), confirming that these organoids resemble the squamous epithelium of the vaginal tissue.

We next sought to generate human vaginal organoids using a human vaginal epithelial cell line called VK2/E6E7 (VK2), which is derived from the vaginal mucosa of a female patient with endometriosis (26). Monolayer and air-liquid interface (ALI) cultures of VK2 cells are susceptible to HSV-2 infection and have been used to interrogate the mechanism of HSV-2 infection *in vitro* (27, 28). These cells also express multiple cytokeratins, which are apical differentiation markers found in the stratified squamous epithelium (26). To generate spheroids, VK2 cells were resuspended in basement membrane extract (BME) and cultured in VK2 media supplemented with TGFβ receptor kinase inhibitor and ROCK inhibitor. By 7 to 10 days post-culture, VK2 cells had aggregated to form spherical organoid-like structures and displayed characteristics of basal cell differentiation, albeit to a lesser degree than mouse vaginal organoids (Fig. 1D). The outer cells of the spheroids expressed both Ki67 and p63, and the inner cells expressed the apical marker filaggrin, showing a similar basal-out structure as the mouse organoids. These data show that human vaginal epithelial cells are also capable of forming differentiated vaginal epithelial spheroids.

### Vaginal organoids and spheroids are susceptible to HSV-2 infection and antiviral drug treatment

To assess whether mouse vaginal organoids and VK2 spheroids support viral replication, we first infected 10 to 14-day old mouse organoids with 2 x 10^6^ PFU/ml of HSV-2 at an approximate multiplicity of infection (MOI) of 1, as determined by cell count at the time of infection. HSV-2 titers were quantified in the supernatants, and the corresponding organoids were used for qRT-PCR and immunofluorescence assays. HSV-2 titer in the supernatants increased significantly at 24 and 48 hours post-infection (Fig. 2A). We also observed increased expression of HSV-2 late genes gC and gE that are involved in complement evasion and viral spread, respectively, as well as the early gene UL9, which is essential for viral replication (Fig. 2B). However, the increase was only significant for gE and UL9. Based on immunofluorescence microscopy, HSV-2 gC protein clustered around the nuclei in basal and suprabasal cells of organoids (Fig. 2C, white arrows). Together, the data show that HSV-2 infects and replicates in mouse vaginal organoids, confirming previous findings by others.

**Fig 2.**
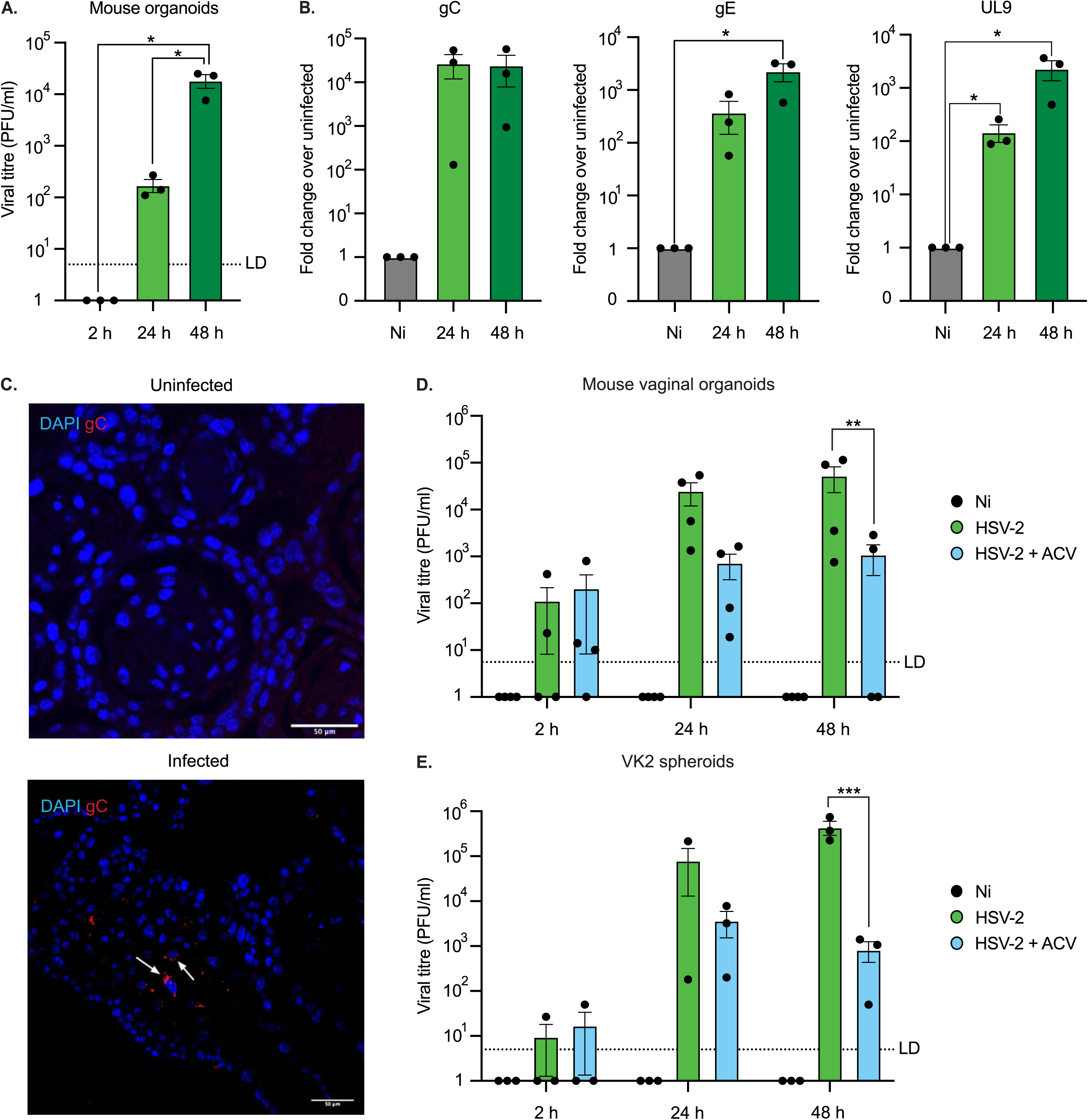
Vaginal organoids and VK2 spheroids are susceptible to HSV-2 infection and responsive to acyclovir treatment. HSV-2 viral titer in the supernatant (A) and viral transcript expression (B) in mouse vaginal organoids infected with HSV-2 at MOI of 1. (C) HSV-2 gC staining on organoid sections at 48 hours post-infection. HSV-2 titer of mouse vaginal organoids (D) and VK2 spheroids (E) infected with HSV-2 in the presence of 200 µM ACV and viral titer. Values are mean ± SEM of 3 biological replicates. For viral gene expression, one-tailed Students’ t-tests comparing Ni with 24 h or 48 h. For all other experiments, one-way ANOVA or two-way ANOVA with Tukey’s test for multiple comparisons. *, p < 0.05; **, p < 0.01; ***, p < 0.001. Magnification 63X; scale bars represent 50 µm. LD, limit of detection (5 PFU/ml). h, hours post-infection.

Next, we investigated whether HSV-2 replication in mouse vaginal organoids could be restricted by acyclovir, an inhibitor of viral DNA replication that has been used to treat HSV infections for decades (29). Organoids were infected with 2 x 10^6^ PFU/ml of HSV-2 in the presence or absence of 200 µM acyclovir (ACV), and viral replication was quantified using plaque assay. HSV-2 replication was significantly reduced at 48 hours post-infection in ACV-treated organoids compared to untreated organoids (Fig. 2D). Similarly, when VK2 spheroids were infected with HSV-2 in the presence or absence of ACV, HSV-2 replication was inhibited by ACV treatment, especially at 48 hours post-infection (Fig. 2E). We also observed comparable levels of HSV-2 replication between VK2 spheroids and mouse organoids. These data show that vaginal organoids and spheroids can also be used to study the effects of antiviral drugs on viral replication.

### ZIKV replicates in mouse vaginal organoids and VK2 spheroids, but drug sensitivity differs between species

Although ZIKV is primarily a mosquito-borne pathogen, sexual contact is also a mode of ZIKV acquisition and transmission (11). Sexually transmitted ZIKV infection has been modeled in mice and macaques that are infected intravaginally, which results in high levels of viral replication (25, 30–33). VK2 cells, as a monolayer culture, are susceptible to ZIKV infection and have been used to study vaginal cell infection by ZIKV (34, 35). Though vaginal organoids have not been explored, ZIKV can infect brain organoids, reflecting the neurotropism of the virus (36, 37). Based on the success of HSV-2 infection in both vaginal organoids and spheroids, we sought to assess ZIKV replication in these models.

Mouse vaginal organoids at 10 to 14-days post-culture were infected with 2 x 10^6^ PFU/ml of ZIKV at an approximate MOI of 1. Viral replication was assessed using enzyme-linked immunosorbent assay (ELISA) for secreted non-structural protein 1 (NS1), which is essential for ZIKV replication. NS1 was detected at 24, 48 and 72 hours post-infection (Fig. 3A). We confirmed ZIKV replication by quantifying the expression of the viral RNA polymerase NS5 by qRT-PCR. The expression of NS5 increased significantly at 48 hours post-infection compared to uninfected organoids (Fig. 3B). We also measured expression of the immune evasion protein NS4B by immunofluorescence microscopy and found that NS4B was detectable in multiple layers of the epithelium in the infected organoids (Fig. 3C, white arrows), but not in uninfected organoids. These findings demonstrate that mouse vaginal organoids can also sustain ZIKV replication.

**Fig 3.**
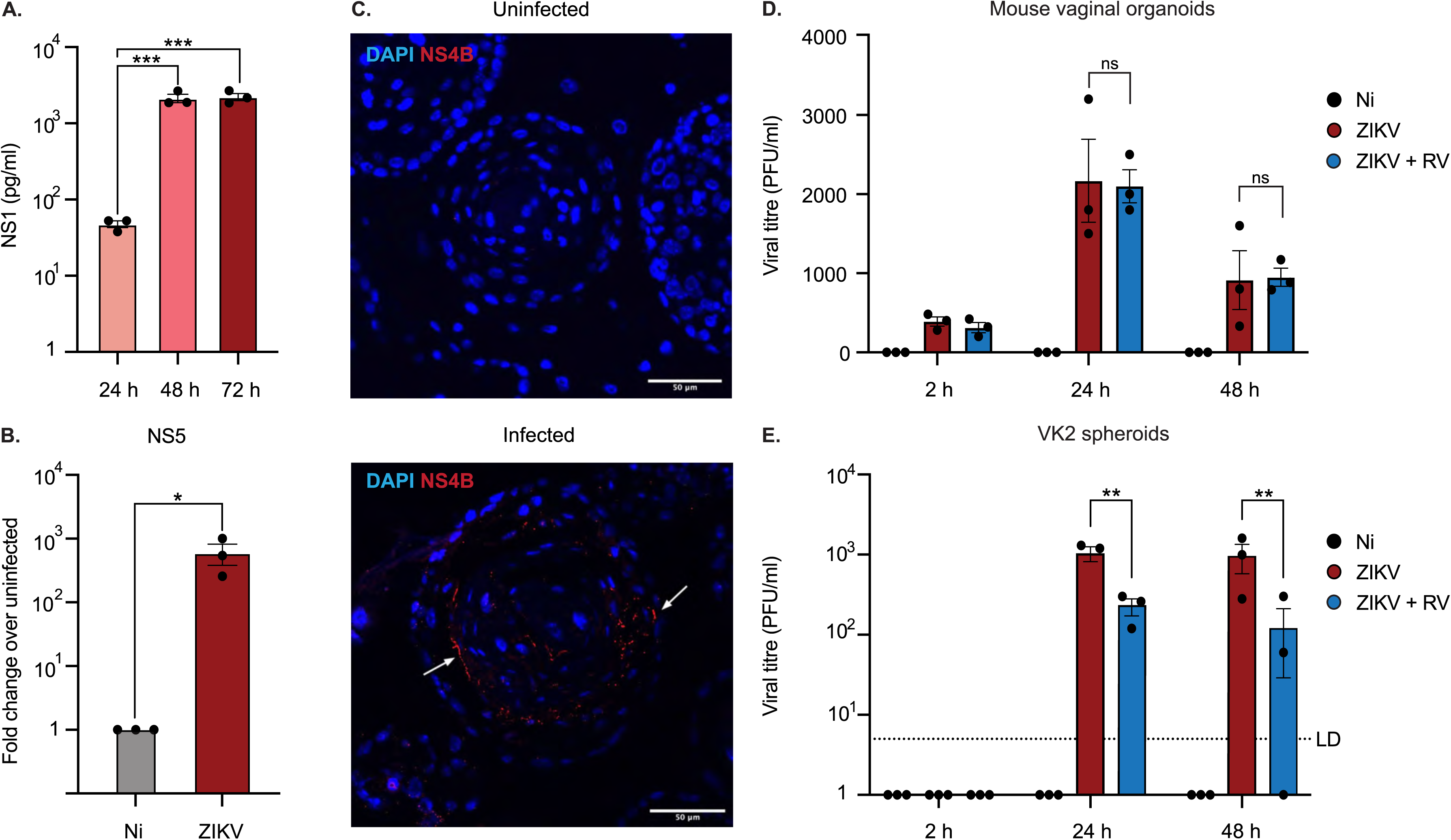
ZIKV replicates in vaginal organoids and VK2 spheroids and is restricted upon remdesivir treatment in VK2 spheroids. NS1 concentration in the supernatant (A) and NS5 transcript expression (B) in mouse vaginal organoids infected with ZIKV at MOI of 1. Values normalized to uninfected samples (Ni) and reported as fold change. (C) NS4B staining in organoids sections from 48 hours post-infection. ZIKV titer of mouse vaginal organoids (D) and VK2 spheroids (E) infected with ZIKV in the presence of 10 µM RDV. Values are mean ± SEM of 3 biological replicates. One-way ANOVA or two-way ANOVA with Tukey’s test for multiple comparisons was performed for statistical analysis. *, p < 0.05; **, p < 0.01; ***, p < 0.001. Magnification 63X; scale bars represent 50 µm. LD, limit of detection (5 PFU/ml). h, hours post-infection.

In contrast to HSV-2, there are no clinically approved antivirals to treat ZIKV infection (38). Remdesivir (RDV), an FDA-approved antiviral drug to treat Ebola and SARS-CoV-2 infections (39) inhibits ZIKV replication *in vitro* (40), but its effect in vaginal cells has never been tested. Thus, we infected both mouse vaginal organoids and VK2 spheroids with 2 x 10^6^ PFU/ml of ZIKV in the presence or absence of 10 µM RDV and quantified viral titer using a plaque assay. In mouse vaginal organoids, ZIKV was measurable at 24 and 48 hours post-infection, but there was no effect of RDV on ZIKV replication (Fig. 3D). ZIKV replication was similarly detectable in VK2 spheroids at 24 and 48 hours post-infection, but in contrast to mouse organoids, RDV treatment resulted in a significant decrease in ZIKV replication (Fig. 3E), revealing species differences in drug efficacy using these models.

### Mouse vaginal organoids have an intact type I IFN response

Type I IFNs are potent antiviral effectors that serve protective roles during vaginal HSV-2 infection (41). Type I IFNs bind to the IFN α/β receptor (IFNAR), resulting in the expression of interferon stimulated genes (ISGs) that further limit viral replication (42). IFN responsiveness varies between differentiation states of cells (43), whereby stem cells are less responsive than differentiated cells. Thus, we sought to assess whether vaginal organoids that arise from the differentiation of the stem cell niche in vaginal tissues are responsive to IFN. To test this, mouse vaginal organoids were treated with 50 ng or 100 ng of recombinant mouse IFNα or infected with 2 x 10^6^ PFU/ml or 2 x 10^7^ PFU/ml of HSV-2 for 6, 12, or 24 hours, and ISG expression was measured by qRT-PCR. The expression of *Mx1*, *Isg15*, *Rsad2*, and *Ifit3* was significantly elevated at 12 hours and 24 hours post-IFNα treatment compared to untreated controls (Fig. 4A-D). At 6 hours post-IFNα treatment, the expression of *Rsad2* and *Ifit3* was also significantly elevated (Fig. 4C, D). Organoids infected with HSV-2 showed a modest increase in the expression of all four ISGs compared to uninfected organoids at all timepoints, albeit not significant. These results show that mouse vaginal organoids can respond to IFN treatment and sense viral infection, indicating that they have an intact type I IFN response.

**Fig 4.**
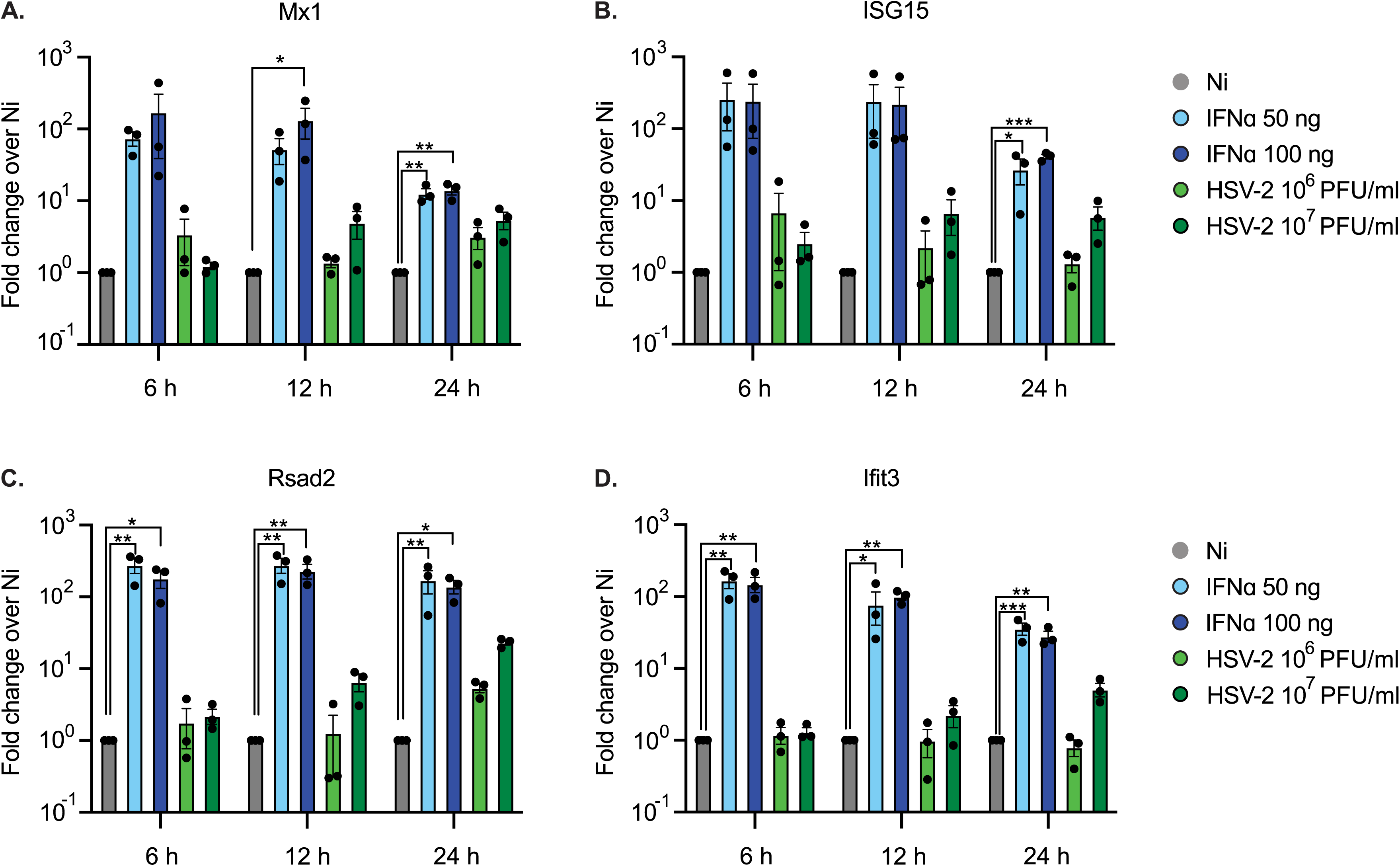
Mouse vaginal organoids express ISGs in response to IFN treatment and HSV-2 infection. (A-D) ISG expression in organoids treated with 50 ng or 100 ng IFNα or infected with HSV-2 at MOI of 1 or 10. All values normalized to uninfected samples (Ni) and presented as fold change. Values are mean ± SEM of 3 biological replicates. One-way ANOVA with Dunnett’s test for multiple comparisons was performed at each time point. *, p < 0.05; **, p < 0.01; ***, p < 0.001. h, hours post-treatment.

### Characterization of apical-out mouse vaginal organoids infected with HSV-2 and ZIKV

Mouse vaginal organoids used in this study and previous studies display a basal-out orientation, whereby basal cells are on the outer surface, while apical cells are at the centre of the organoids facing the lumen. The basal-out orientation commonly develops when organoids are cultured in the presence of Matrigel or BME, substances which are analogous to the extracellular matrix (ECM). While basal-out organoids are susceptible to HSV-2 and ZIKV infection, they may not fully recapitulate primary infection through the apical surface.

Previous studies have shown that basal-out organoids can be flipped into apical-out organoids when they are released from Matrigel or BME and cultured in suspension (44). We tested this by dissociating 6 to 7 day old basal-out mouse vaginal organoids from BME and culturing them in suspension (Fig. 5A). On day 1 post-suspension culture, organoids retained the basal-out orientation, but they started to flip on day 3 and fully formed apical-out organoids by day 5 post-culture (Fig. 5B). After 10 days of culture in suspension, these organoids were fixed and sectioned. H&E staining revealed the presence of a large lumen surrounded by 2-3 layers of cells (Fig. 5C). Staining for filaggrin, p63, and Ki67 further showed that the basal cell markers p63 and Ki67 were expressed by cells surrounding the lumen, while the apical cell marker filaggrin was expressed by cells on the external side. Moreover, apical-out organoids expressed cytokeratin 13 on the outer cells, distinct from the Ki67-expressing cells that face the lumen (Fig. 5D). This is contrast to the basal-out organoids, which have a reversed orientation. These studies demonstrate, for the first time, the establishment of apical-out mouse vaginal organoids.

**Fig 5.**
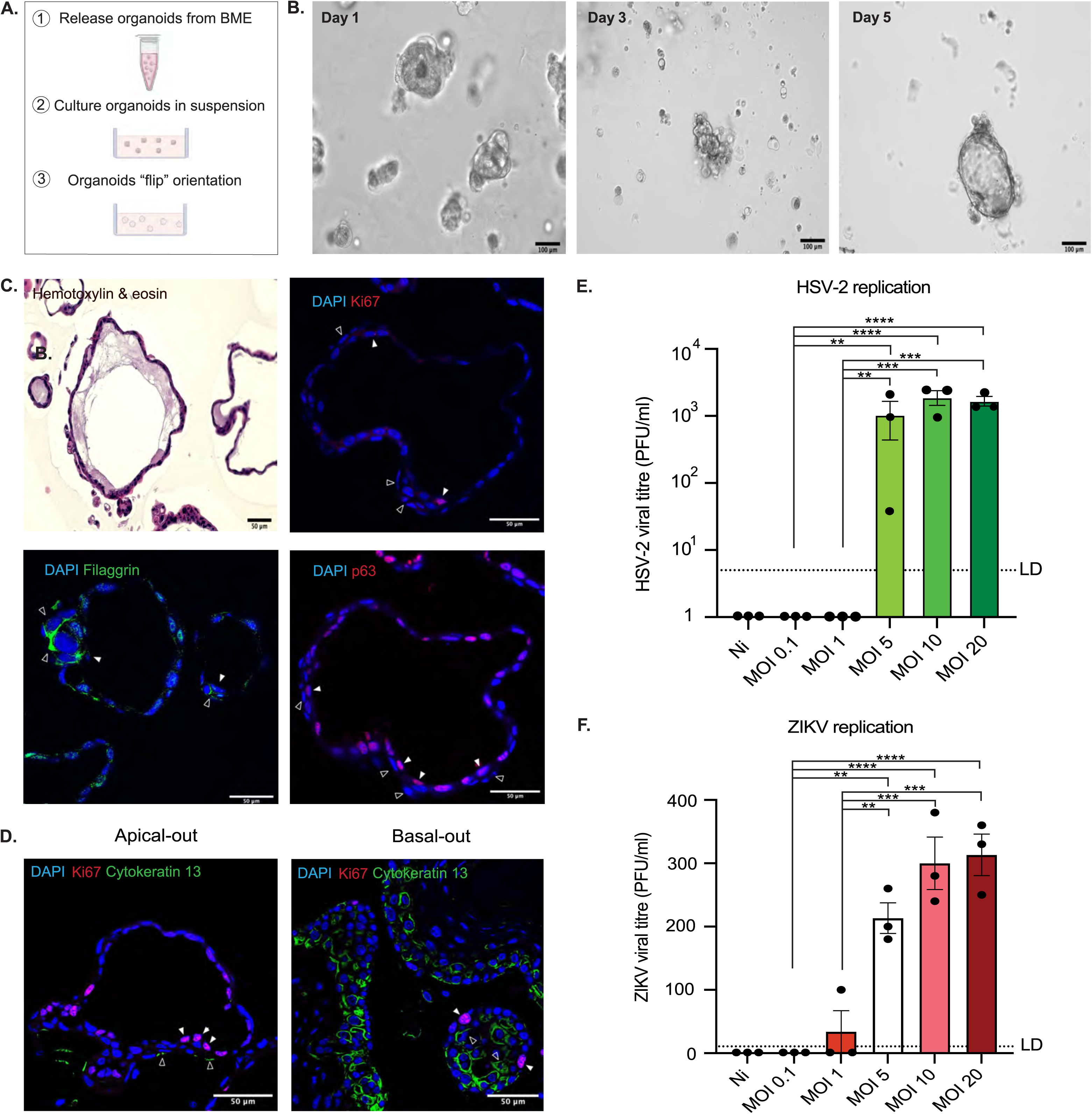
Apical-out mouse vaginal organoids are susceptible to HSV-2 and ZIKV infection. (A) A schematic illustration of the establishment of apical-out vaginal organoids. (B) Brightfield images of apical-out organoid development. (C) Sections of 10 to14 day old apical-out organoids stained for H&E, Ki67, p63, and filaggrin. (D) Comparison of cytokeratin-13 localization between apical-out (left) and basal-out (right) organoids. Viral titers of apical-out organoids infected with HSV-2 (E) and ZIKV (F) at the indicated MOIs for 48 hours. Values are mean ± SEM of 3 biological replicates. Two-way ANOVA with Tukey’s test for multiple comparisons was performed for statistical analysis. *, p < 0.05; **, p < 0.01; ***, p < 0.001; ****, p < 0.0001. Magnification 20X (brightfield) and 63X; scale bars represent 50 µm. LD, limit of detection (5 PFU/ml). h; hours post-infection. Solid arrows indicate basal cells and open arrows indicate apical cells.

To determine whether apical-out mouse vaginal organoids are susceptible to HSV-2 and ZIKV infection, we infected 7 day-old apical-out organoids with HSV-2 and ZIKV at MOI of 0.1, 1, 5, 10, and 20 and measured viral replication at 48 hours post-infection. In contrast to the basal-out organoids, apical-out organoids required a much higher MOI for efficient HSV-2 replication (Fig. 5E). Similarly, significant replication of ZIKV was observed upon infection at MOI of 5 or higher (Fig. 5F). Overall, these results indicate that apical-out organoids support viral replication but require a higher amount of virus to support productive infection compared to basal-out organoids.

## Discussion

Our study is the first to demonstrate that acyclovir blocks HSV-2 replication in mouse and human vaginal organoids. Many novel anti-herpetic drugs are currently under development, including non-nucleoside kinase inhibitors, topical interferon formulations, and virucidal spirocyclic compounds, which could combat rising rates of acyclovir-resistant HSV-2 infections (45–48). Many of these drug candidates are initially screened in non-human primate Vero cells (45, 46, 48), which might not account for species-specific or cell type-specific effects on antiviral activity. The mouse and human vaginal organoids described in this study are useful alternatives for screening novel drug candidates, as they maintain the structural complexity of the vaginal epithelium and are scalable for high-throughput drug screens (49).

Although species differences have been reported for the ability of ZIKV to antagonize the IFN response (50), our data showed similar levels of ZIKV replication in both mouse and human vaginal organoids, supporting previous *in vivo* studies of intravaginal ZIKV infection in mice (30–32). Our results in organoids confirmed previous findings that ZIKV is able to replicate in mouse vaginal tissues derived from wildtype IFN-competent mice (25, 50). However, we found species differences in the effect of RDV against ZIKV replication. A possible explanation for this could be due to differences in RDV metabolism, as has been shown when comparing non-human primate cells to human cells (51, 52). More studies are needed to extend RDV testing in vaginal cells and to better understand species differences in drug metabolism in the vaginal tract.

Type I IFN signalling is an important antiviral response (53), and mice lacking IFN signalling are highly susceptible to vaginal HSV-2 infection (54). Mouse vaginal organoids showed robust ISG expression in response to IFNα treatment, indicating that, as expected, the IFNAR signalling pathway remains intact even after redifferentiation of vaginal stem cells into differentiated epithelial cells. HSV-2 infection also resulted in ISG expression in the mouse vaginal organoids, albeit modestly. This likely reflects the ability of HSV-2 ICP27 to antagonize the IFN response via IRF3 (27, 55). Our data indicate that mouse vaginal organoids can serve as useful tools to probe innate immune responses to STVs.

The basal-out orientation of vaginal organoids limits access to the apical surface and could affect viral infection. To better assess how this orientation affects viral infection, we generated apical-out mouse vaginal organoids by culturing cells in suspension instead of in BME. Apical-out organoids exhibited notable structural differences compared to basal-out organoids, such as fewer layers of epithelial cells and the presence of a large central cavity, which has been observed in intestinal organoids (44, 56). These differences can be explained in part by the physical constraints of the ECM domes, as the presence or absence of ECM has been shown to affect lumen formation in organoid cultures. Cellular aggregates grown in the presence of Matrigel form lumens by membrane restructuring and separation through a process called hollowing, whereas those grown without Matrigel form lumens via apoptosis of central cells through a process called cavitation (57). Future investigations into the effects of these structural differences in the epithelium and lumen formation on viral infection may yield novel insights into host-pathogen interactions.

HSV-2 infection of apical-out organoids revealed that these organoids required a higher MOI for productive replication compared to basal-out organoids. This difference was unexpected given that HSV-2 entry receptors Nectin-1 and HVEM are expressed at the apical surface of the vaginal epithelium (58, 59). Previous studies have shown that Nectin-1 expression in the murine vaginal epithelium varies during the estrous cycle and across epithelial layers (58, 60, 61). Nectin-1 expression and HSV-2 susceptibility are higher when estradiol levels are elevated. Although our mice were treated with DMPA and synched in diestrus phase, our culture media lacked estrogen, which might have reduced the level of Nectin-1 expression and decreased HSV-2 susceptibility in apical-out organoids. Alternatively, another possible explanation for the difference in viral susceptibility is that apical cells are better able to restrict viral replication compared to basal cells. It has been previously shown in human and murine nasal and skin epithelia that IFN and ISG expression differs across epithelial cell layers, with higher expression in apical cells compared to basal cells (62, 63). Although this has not yet been studied in the vaginal epithelium, it is plausible that apical vaginal epithelial cells are better at restricting viral infection due to increased IFN expression. Altogether, these possibilities highlight key features of the vaginal epithelium that are important to consider for future work using vaginal organoid models.

A previous study established 3D aggregates of the immortalized human vaginal cell line VI9I using a rotating wall vessel (RWV) bioreactor and showed that these aggregates express markers of apical differentiation (64). In our study, we showed that BME was sufficient to generate 3D spheroids from a human vaginal cell line without the need for a RWV bioreactor. These spheroids expressed both apical and basal cell markers and were susceptible to infection by HSV-2 and ZIKV. To the best of our knowledge, this is the first time that 3D cultures of human vaginal epithelial cells have been used to model viral infection and test the effect of antiviral drugs, providing a useful tool for future studies on human vaginal infections and viral pathogenesis.

In summary, we demonstrated that mouse and human vaginal organoid systems are capable of sustaining viral replication and responding to antiviral drugs. These results contribute to the ongoing characterization of these recently established organoids and provide additional evidence of their ability to serve as models of vaginal viral infections.

## Materials and Methods

### Mice

C57BL/6N (B6) mice were bred in the Modified Barrier Facility at UBC. 10- to 12-week-old female mice were used to establish vaginal organoids cultures. All procedures performed complied with the policies set by the UBC Animal Care Committee (UBC A21-0093).

### Viruses

HSV-2 strain 333 is previously described (63) and was a kind gift from Dr. Marc Horwitz at the University of British Columbia. ZIKV strain PRVABC59 was a kind gift from Drs. Selena Sagan and Francois Jean at the University of British Columbia. HSV-2 GFP was a kind gift from Dr. Ali Ashkar at McMaster University, Hamilton, Ontario.

### Viral propagation and quantification

HSV-2 was propagated on Vero cells. Briefly, Vero cells were infected at MOI 0.05 in 10 ml plain DMEM for 1 hour at 37 °C. Media was replaced with DMEM containing 1% FBS and 1% Penicillin-Streptomycin (PS) and incubated at 37 °C for 2 days. After 2 days, infected cells and supernatant were collected in 2.5 ml aliquots and an equal volume of autoclaved 0.3 M skim milk was added to each aliquot before freezing overnight at -80 °C. Aliquots were thawed and infected cells were sonicated on ice as follows: 30 seconds on, 30 seconds off for 3 cycles at maximum amplitude. Sonicated material was clarified by centrifuging at 1600 rpm for 10 minutes at 4 °C. Supernatant was collected and stored at -80 °C.

HSV-2 titer was quantified using plaque assay. Briefly, Vero E6 cells were plated in 6 well plates at 0.37 x 10^6^ cells per well. The next day, supernatants were thawed on ice and diluted 10fold in plain DMEM. Vero cells were infected with 350 µl/well of diluted virus in plain DMEM. Plates were incubated at 37°C, 5% CO_2_ for 1 hour and rocked every 20 minutes. After one hour, viral inoculum was aspirated and replaced with 2 ml per well of DMEM supplemented with 1% FBS, 1% PS, and 2 mg/ml human IgG (Sigma Aldrich Cat No. I4506-10MG). Cells were incubated for 2 days and fixed in methanol with 0.1% crystal violet.

ZIKV was passaged in Vero cells. Briefly, 10^5^ Vero cells were seeded in each well of a 6-well plate and were infected with ZIKV at MOI 0.01 in Eagle’s Minimum Essential Medium (EMEM). After incubating at 37 °C for 2 hours, media was replaced with DMEM (2% FBS, 1% PS, 1% L-glutamine, 1% NEAA, and 15 mM HEPES), and cells were incubated at 37°C, 5% CO_2_ for 3 days. Supernatants were clarified by centrifugation at 300 g for 10 minutes and stored at -80 °C. The resulting titer of the passaged virus was quantified by plaque assay on Vero cells, as described previously (64).

ZIKV titer from infection experiments was quantified using an ELISA to detect NS1 concentration (Sino Biological, Cat No. KIT40544). ELISAs were conducted according to manufacturer’s instructions and absorbance data was graphed using Microsoft Excel.

### Primary mouse vaginal organoid cultures

*In vitro* cultures of primary mouse vaginal organoids were established as described previously (23). Briefly, 10- to 12-week-old female B6 mice were injected subcutaneously with 20 mg/ml DMPA (Pfizer), and vaginal tissues were harvested on day 5. Tissues were slit to expose the epithelium and digested in DMEM/F12 containing 1 mg/ml Pronase (Sigma Cat No. 10165921001), 0.5 mg/ml DNaseI (Sigma Cat No. 10104159001), and P/S (Gibco Cat No. 15140122) at 4°C for 15 hours with gentle shaking. Enzymatic digestion was stopped by adding DMEM/F12 (Thermo Cat No. 11320033) containing 10% FBS. Vaginal tissues were minced with scissors, centrifuged at 200 g for 5 min at 4°C, washed twice in DMEM/F12, and filtered through a 40 µm cell strainer. Cells were pelleted by centrifuging at 200 g for 5 min at 4°C, resuspended in DMEM/F12 containing FBS. For organoid cultures, 20,000 cells were resuspended in 50 µl BME (R&D Systems Cat No. 3533-010-02) and cultured in DMEM/F12 supplemented with 2% Ultroser G (Sartorius Cat No. 15950-017), 1% PS, 50 ng/ml mouse EGF (Sigma Cat No. SRP3196-500UG), 0.5 mM SB431542 (Selleck Chemicals Cat No. S1067), and 10 mM Y-27632 (Tocris Cat No. 1254) (OCM). Medium was changed every two days, and Y27632 was removed after three days.

### VK2 spheroids

VK2 cells were a kind gift from Dr. Charu Kaushic at McMaster University, Hamilton, Canada. Cells were grown as previously described (65) and cultured in complete keratinocyte serum free media (KSFM; Gibco Cat No. 17005042). For spheroid cultures, cells were rinsed with 0.25% trypsin-0.03% EDTA solution, trypsinized with the trypsin-EDTA solution for 10 min at 37 °C, 5% CO_2_. Trypsin was neutralized using DMEM/F12 containing 10% FBS. Cells were centrifuged at 120 x g for 10 min and resuspended in KSFM. 20,000 cells were resuspended in 50 µl BME and cultured in KSFM supplemented with 1% PS, 0.5 mM SB431542, and 10 mM Y-27632. The medium was changed every two days, and Y-27632 was removed after three days.

### *In vitro* culture of apical-out mouse vaginal organoids

*In vitro* cultures of apical-out mouse vaginal organoids were established from cultures of basal-out mouse vaginal organoids. 5 to 7 day old basal-out organoids were released from BME by forcefully adding 1 ml of ice-cold DPBS (Invitrogen Cat No. J67802.AP) per well and pipetting up and down 20 times. Organoids were collected and centrifuged at 1100 rpm for 5 minutes at 4°C. Supernatants containing DPBS and BME shards were aspirated, and organoids were resuspended in DPBS. This washing step was repeated 4 to 5 times. After the last wash, organoids were resuspended in OCM without Y-27632, and 500 µl of the suspension was transferred to a 24-well Ultra-Low attachment plate (Corning Cat No. 3473). The conversion of basal-out organoids to apical-out organoids was observed over 3 to 5 days post-culture using a EVOS M3000 (Thermo) microscope and a 20X objective.

### Viral infection and IFNα treatment of organoids

To calculate the MOI, organoids were dissociated and cells were counted. The average number of cells from 3 wells was used to calculate the MOI. Media was removed from organoids and replaced with 500 µl/well of virus diluted in pre-warmed DMEM (Sigma Aldrich Cat No. D5671). Cells were incubated at 37°C, 5% CO_2_ for 2 hours, then viral inoculum was removed, and organoids were washed thrice with pre-warmed DMEM to remove the inoculum. Organoid culture media was added to wells, and cells were incubated at 37°C, 5% CO_2_. At the indicated time points, supernatants were collected, and organoids were harvested for subsequent RNA extraction.

For drug experiments, basal-out vaginal organoids were infected with HSV-2 and ZIKV for 2 hours as described above to allow for viral entry. After 2 hours, viral inoculum was removed and organoids were washed in plain DMEM 4-5 times. The last wash was collected as the 2 hour time point, and 500 µl of OCM supplemented with 200 µM Acyclovir or 10 µM Remdesivir. Supernatants were collected at 24 and 48 hours post-infection, and viral titer was quantified as described above.

For ISG expression experiments, basal-out organoids were infected with HSV-2 or treated 50 ng or 100 ng of mouse interferon ɑ (Thermo Cat No. 14831280) in 500 ul of pre-warmed OCM. Viral inoculum was kept in the culture for the duration of the experiment.

For infection of apical-out organoids with HSV-2 GFP, 7 day old organoids were infected for 2 hours with 500ul of 2 x 10^6^ PFU/ml of HSV-2 GFP or with HSV-2 and ZIKV at MOI of 0.1, 1, 5, 10 and 20 in plain DMEM. After 2 hours, organoids were harvested and centrifuged at 1100 rpm for 5 minutes. Supernatant was aspirated and organoids were resuspended in 500 µl plain DMEM. The washing step was repeated 5 times. Finally, organoids were resuspended in 500 µl OCM and returned to a 24 well ultra-low attachment plate. For HSV-2 GFP infection, organoids were visualized on day 5 post-infection on the EVOS M3000 (Thermo) using a 10X objective. For HSV-2 and ZIKV infection, supernatants were collected at 48 hours post-infection after centrifuging organoids at 300 g for 5 minutes at 4°C.

### Immunofluorescence confocal microscopy of vaginal organoids, vaginal tissue, and VK2 spheroids

BME domes of mouse organoid cultures and spheroids were disrupted by forcefully adding 1 ml of ice-cold DPBS per well. Organoids were released from BME by pipetting up and down 20 times, collected and fixed in 4% PFA for 30 min at room temperature. Vaginal tissues were fixed in 4% PFA for 30 min at room temperature. Fixed organoids and vaginal tissues were washed three times in PBS for 10 min each and embedded in 2% agarose, followed by paraffin fixation, embedding, and sectioning at 4µm for hematoxylin & eosin staining (ACS Research Histology Laboratory at UBC).

For immunofluorescence staining, paraffin-embedded sections of vaginal organoids and vaginal tissues were deparaffinized in xylene and rehydrated through descending grades of ethanol. Antigens were unmasked by heating at 95°C for 30 minutes in 1mM EDTA pH 8.0 or 1X Antigen Retrieval Solution (Dako Cat No. S1699). Slides were rinsed thrice with PBS, then blocked for 1 hour at room temperature in PBS with 0.1% Triton-X100 and 1% normal goat serum (Thermo Fisher). Sections were incubated overnight with primary antibodies at 4°C, then washed thrice with 0.1 % Tween-20 in PBS and incubated with secondary antibodies for 2 hours at room temperature. Slides were rinsed with 0.1% Tween-20, incubated with Hoescht-33342 (1:500) for 15 minutes at room temperature, rinsed with 0.1% Tween-20, and mounted in ProLong Gold Antifade (Thermo Fisher Cat No. P36935). For co-staining with apical and basal markers, slides were incubated overnight with primary antibodies against apical markers at 4°C, then washed thrice with 0.1% Tween-20 in PBS and incubated with secondary antibodies and directly conjugated primary antibody against basal markers for 2 hours at room temperature. For Nectin-1 staining of vaginal organoid sections, deparaffinization, rehydration, and antigen retrieval was performed as above. Sections were blocked using an endogenous biotin-blocking kit (Invitrogen, Cat. No. E213390) and staining was performed using the Vector M.O.M. Immunodetection kit (Vector, Cat. No. FMK-2201) according to manufacturer’s instructions.

For whole mount staining of vaginal organoids, fixed organoids were washed thrice in 100 mM glycine in PBS, then blocked and permeabilized overnight at room temperature in PBS with 0.3% Triton X-100 and 5% normal goat serum (Thermo Scientific Cat No. 01-6201). Primary antibodies in PBS containing 0.3% Triton X-100 and 1% normal goat serum were added overnight, followed by addition of secondary antibodies overnight. Hoechst-33342 was added at a final dilution of 1:500 and incubated for 15 minutes at room temperature. Organoids were washed once with dH_2_O, allowed to settle for 5 minutes, washed once with PBS, and incubated for 1 hour each in 50% and 100% methanol in the dark at room temperature. Methanol was removed, organoids were transferred to chamber slides in ProLong Gold Antifade Mountant and dried overnight before imaging.

The following primary antibodies were used: Ki67-FITC (Rat, Thermo Fisher, Cat. No. 11-5698-82), Filaggrin (Rabbit, BioLegend, Cat. No. 905804), p63 (Rabbit, Abcam, Cat. No. EPR5701), E-cadherin (Rat, Thermo Fisher, Cat. No. 53-3249-82), Nectin-1 (Mouse, Invitrogen, Cat. No. 37-5900), HSV-2 gC (Mouse, SantaCruz, Cat. No. sc69801), and ZIKV NS4B (Rabbit, GeneTex, Cat. No. GTX133321). The following secondary antibodies were used: Goat anti-mouse AlexaFluor488 (Invitrogen, Cat. No. A28175), donkey anti-rabbit AlexaFluor680 (Invitrogen, Cat. No. A10043), and goat antirabbit Cy3 (Jackson, Cat. No. 111-165-003). Images were obtained on a Leica SP5 confocal microscope containing LAS X software using a 63X oil immersion objective. Images were exported as TIFF files and analyzed using FIJI software.

### RNA extraction and qRT-PCR

Media was aspirated and BME drops were disrupted by forcefully adding 1 ml of PBS containing 1% BSA. Organoids were released from BME by pipetting up and down 20 times, collected in tubes, and centrifuged at 300 x g for 10 min at room temperature. Organoid pellets were resuspended in 300 ul of RNA lysis buffer (RNeasy mini kit, Qiagen Cat No. 74104), and RNA was extracted according to the manufacturer’s instructions. 1 ug of RNA was reverse transcribed to cDNA using the iScript cDNA synthesis kit (BioRad Cat No. 1708891). RT-qPCR was performed using QuantStudio 3 (Thermo Fisher) and PowerTrack SYBR Green Master Mix (Thermo Fisher Cat No. A46110). The following amplification settings were used: 95°C for 2 minutes, followed by 40 cycles of 95°C for 15 seconds and 60°C for 1 minute. Melt curve analysis was conducted at 95°C for 15 seconds, 60°C for 1 minute, and 95°C for 1 second. Fold change was calculated using the ΔΔCt method. The following primers were used:

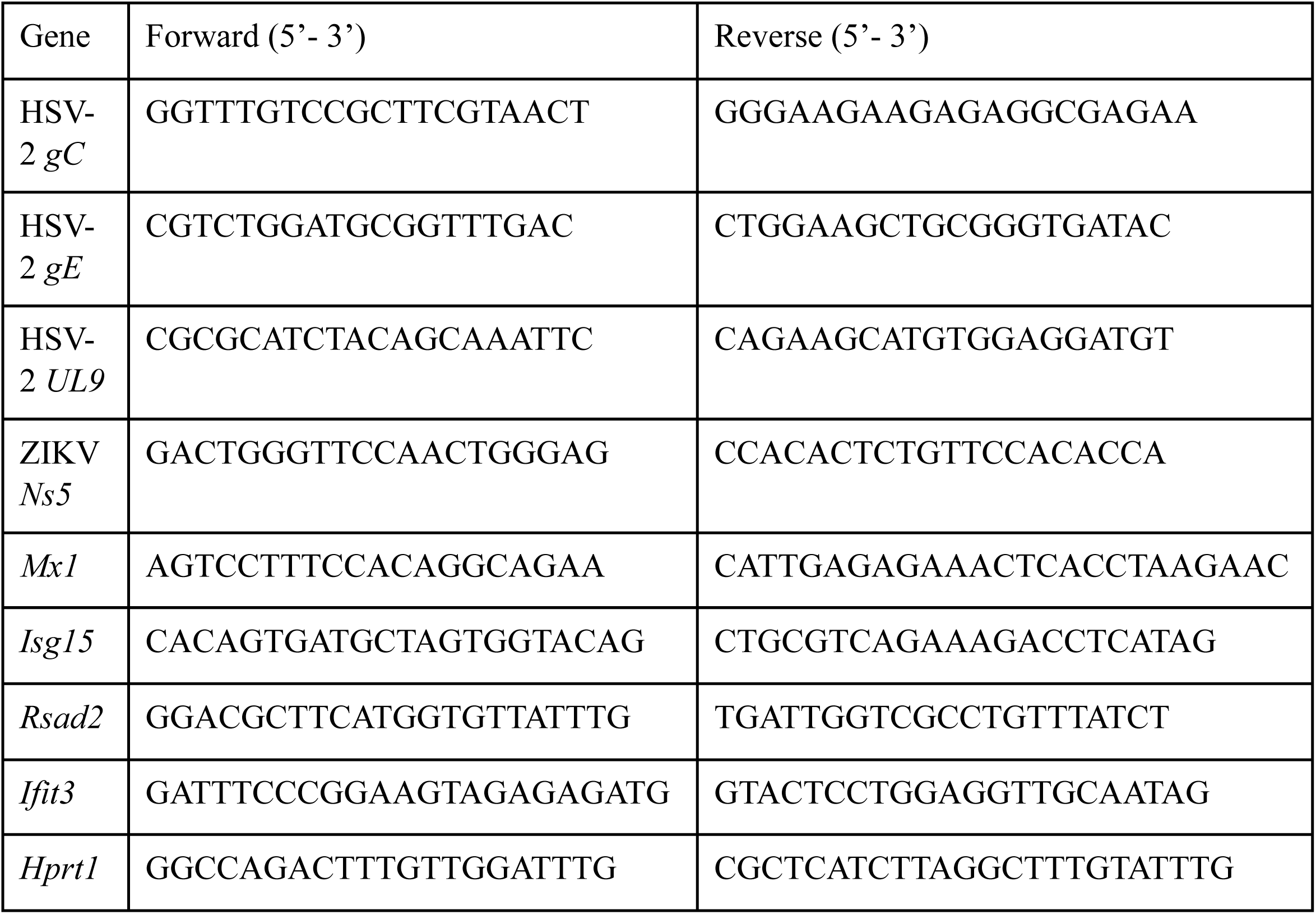

## Acknowledgements

This work was supported by the Canadian Institutes of Health Research (PJT 183573 and ECP184181 to M.T.), Michael Smith Health Research Scholar Award (SCH-2022-2804 to M.T.), and a University of British Columbia Affiliated Master’s Fellowship (to A.S.). We would like to thank Dr. Janelle Kopp, Kevin Wang, and Dr. Leah Hohman for assisting with the immunofluorescence assays. We acknowledge the Dr. Guang Gao at the UBC High Resolution Microscopy Core and the UBC Modified Barrier Facility for facilitating the microscopy and animal experiments, respectively. Figures were generated using BioRender. We also acknowledge that this work was conceptualized and carried out on the traditional, ancestral, and unceded territory of the xʷməθkʷəy̓ əm (Musqueam) people.

## Author Contributions

Conceptualization, A.S. and M. T.; experimentation, A.S., I.P., G.B., I.B., and M.D.L.; data analysis, A.S. and I.P; writing, A.S. and M.T.; funding acquisition, M.T.

